# Glycoproteome Analysis of Human Serum and Brain Tissue

**DOI:** 10.1101/647081

**Authors:** Christopher J. Brown, Kathleen T. Grassmyer, Matthew L. MacDonald, David E. Clemmer, Jonathan C. Trinidad

## Abstract

Protein glycosylation represents one of the most common and heterogeneous post-translational modifications (PTMs) in human biology. Herein, an approach for the enrichment of glycopeptides using multi-lectin weak affinity chromatography (M-LWAC), followed by fractionation of the enriched material, and multi-mode fragmentation LC/MS is described. Two fragmentation methods, high-energy collision induced dissociation (HCD) and electron transfer dissociation (EThcD), were independently analyzed. While each fragmentation method provided similar glycopeptide coverage, there was some dependence on the glycoform identity. From these data a total of 7,503 unique glycopeptides belonging to 666 glycoproteins from the combined tissue types, human serum and brain, were identified. Of these, 617 glycopeptides (192 proteins) were found in both tissues; 2,006 glycopeptides (48 proteins) were unique to serum, and 4,880 glycopeptides (426 proteins) were unique to brain tissue. From 379 unique glycoforms, 1,420 unique sites of glycosylation were identified, with an average of four glycans per site. Glycan occurrences were significantly different between tissue types: serum showed greater glycan diversity whereas brain tissue showed a greater abundance of the high mannose family. Glycosylation co-occurrence rates were determined, which enabled us to infer differences in underlying biosynthetic pathways.

## Introduction

Glycosylation, the co- and post-translational attachment of glycans to polypeptides on secreted and membrane regions, is one of the most prevalent PTMs (1). This PTM involves a heterogeneous family of glycans and regulates multiple aspects of protein activity including conformation (2, 3), stability (4) and protein-protein interactions (5). Glycosylation is a hallmark of the immune system, where it regulates activation of one arm of the complement pathway (6). Aberrant protein glycosylation has been implicated in a variety of diseases (7–9), ranging from cancer (10–15) to inflammatory diseases (6, 16–19) to congenital disorders of glycosylation (20, 21). Despite the key roles played by glycosylation, we still have an incomplete understanding of its extent and regulation in either healthy or disease states.

The two most common types of protein glycosylation are O-linked and N-linked, with the latter being more widespread (22). N-linked glycans, the focus of this manuscript, are attached to the nitrogen of asparagine residues, typically at an N-X-S/T motif, where X is any amino acid but proline (22). The principle N-glycan subtypes are high-mannose, complex and hybrid, which may be further modified with structures such as bisecting GlcNAc, fucose, and sialic acid.(23) These additional modifications modulate protein activity with particular implications for cancer (12, 13, 24, 25). Proteins can exist as a variety of glycoforms, where a glycosylation site can be occupied by numerous distinct glycans, a process termed microheterogeneity. Glycosylation can be studied from a glycomics perspective, where glycans are released from glycoproteins allowing examination of global glycan levels (26, 27). Alternatively, techniques exist to identify sites previously glycosylated prior to glycan release (14, 28). However, comparatively fewer large-scale studies addressed the exact identity of glycans modifying specific sites on the proteome to provide a more integrated view (29–33).

Lectin-based separations combined with mass spectrometry (MS) are a powerful means for identifying glycopeptides. Recent work, by Riley et al. identified over 5,600 glycopeptides from murine brain (29). This extends our previous work using immobilized lectins that mapped over 2,500 unique glycopeptides from similar tissue (31). Our initial work relied on a column of wheat germ agglutinin (WGA), which is relatively specific for N-acetyl-D-glucosamine and sialic acid (34). We now investigate a column with six immobilized lectins to more broadly enrich glycopeptides with potentially fewer biases (35–39).

In this paper, we developed an advanced biochemical enrichment and fractionation method to investigate the glycoproteome of human serum and post-mortem brain tissue to better understand the differences in N-glycan processing in these tissues (12). We report the largest human glycoproteome to date with 7,503 total glycopeptides mapping to 666 glycoproteins. Of these, 617 glycopeptides and 192 glycoproteins are found in both tissues. 48 Glycoproteins and 2,006 glycopeptides were only found in serum. 426 Glycoproteins and 4,880 glycopeptides were only found in brain. We determine the extent to which specific glycoforms are more prevalent in a given tissue. We also examine the data from a global perspective to demonstrate how glycoproteomic data can be integrated to assess the underlying glycan synthesis pathways.

## Experimental procedures

### Brain Lysate Sample Preparation

Brain specimens from all subjects were obtained during autopsies conducted at the Allegheny County Office of the Medical Examiner after receiving consent from the next-of-kin. Procedures were approved by the University of Pittsburgh Institutional Review Board and Committee for Oversight of Research Involving the Dead. Grey matter was harvested from the auditory cortex as previously described (40, 41): Tissue slabs containing the superior temporal gyrus with Heschl’s Gyrus located medial to the planum temporale were identified, and the superior temporal gyrus removed as single block. Grey matter (100 mg) was collected from HG by taking 100 μm frozen sections (40). Grey matter was homogenized in 1 mL of 8M urea with a FastPrep-24 benchtop homogenizer. Samples were stored at −20 °C.

### Protein Preparation

Sera (Sigma-Aldrich, St. Louis, MO) was denatured using 8 M urea (Sigma-Aldrich, St. Louis MO) in 100 mM ammonium bicarbonate (*p*H 7.4, Sigma-Aldrich, St. Louis MO). Disulfide bonds were reduced using 1 mM tris(2-carboxyethyl)phosphine (Sigma-Aldrich, St. Louis, MO) for 1 hour at 65 °C. Reduced cysteine residues were modified using 4.4 mM iodoacetamide (Sigma-Aldrich, St. Louis, MO) for 1 hour at room temperature. Following this, the sample was digested overnight at 37 °C with trypsin (1:200 w:w; Promega, Madison, WI). Peptides were desalted using a Sep-Pak (Waters, Milford, MA), dried and stored at –20 °C until use.

Brain lysates were adjusted to 100 mM ammonium bicarbonate and digested as above. Peptides were resolubilized in a buffer containing 100 mM ammonium bicarbonate (*p*H 8.0), 100 mM sodium chloride, and 2 mM calcium chloride and stored at –20 °C until use.

### Multi-lectin Enrichment Column

Lectin functionalized resin was generated using previously described methods (42, 43). POROS resin (ThermoFisher, Waltham, MA) was resuspended in phosphate buffered saline (*p*H 7.5, Sigma-Aldrich, St. Louis MO). Both lectin and NaBH_3_CN (Sigma-Aldrich, St. Louis MO) were added to the POROS resin and mixed overnight at 4 °C. The reaction was quenched through the addition of both 100 mM Tris buffer, and additional NaBH_3_CN, followed by a 4 °C overnight incubation period. The crosslinked resin was then spun down and the supernatant separated to quantify the linkage efficiency. Lectin-functionalized resin was stored in LWAC ammonium biocarbonate (ABC) buffer (100 mM ABC, 50 mM NaCl, 2 mM CaCl_2_, *p*H 8.0) with 0.05% sodium azide. Individual lectins were separately cross linked to the resin. Following cross-linking, the lectin specific resins were mixed in equal amounts and used without further modification. All lectins were acquired from Vector Laboratories (Burlingame, CA) and used without further purification. Lectins used included *Canavalia ensiformis*, *Ulex europaeus*, *Lotus tetragonolobus*, *Sambucus nigra*, *Triticum vulgaris*, and *Artocarpus integrifolia*. Supplemental Materials and Methods Table 1 shows the linkage efficiency of each lectin. The final resin mixture was self-packed using an ÄKTA Pure (GE Biosciences, Marlborough, MA) into a 2 cm × 2.8 mm column, termed the multi-lectin weak affinity chromatography (M-LWAC) column, using ABC LWAC buffer (100 mM ABC, 50 mM NaCl, 2 mM CaCl_2_, *p*H 8.0) with 0.05% sodium azide, and kept refrigerated until use.

### ÄKTA Pure M-LWAC enrichment and fractionation

Five mg of brain digest and 10 mg of sera digest were separately dissolved in 250 µL of ammonium bicarbonate LWAC buffer. 25 µL aliquots were injected onto the ÄKTA pure system using an autosampler, with a LWAC Tris buffer (100mM Tris, 50 mM NaCl, 2mM CaCl_2_, *p*H 7.5) at a flow rate of 200 µL/min. The M-LWAC column was placed in an ice bath during enrichment. Only the M-LWAC column was placed inline initially (Figure 1). At 3.5 minutes after injection, the flow-through peak (previously characterized to contain the bulk of non-glycosylated peptides) was observed to be at 10% maximum intensity (43). At this point the high *p*H C18 trap (self-packed, Poros C18 Resin, Thermo-Fischer, dimensions 2 cm × 2.8 mm.) was placed inline and the LWAC elution was trapped for 90 minutes. This was repeated for a total of 10 injections of 25 µL each, successively trapping all the LWAC eluent. The sample was introduced over 10 injections to not overload the LWAC column. For the 10^th^ injection, after 90 minutes the M-LWAC column was taken offline and the high *p*H C18 trap was placed online with the high *p*H analytical column (Kinetex C18 100Å, 5 µm, 150 mm × 21 mm). Both of these columns were washed at 100 µL/min for 118 minutes with 20 mM ammonium formate (*p*H 10) in water for desalting. The high *p*H reverse phase analytical column utilized this aqueous buffer (A) and 20mM ammonium formate in 80% acetonitrile (ThermoFisher, Waltham, MA) at *p*H 10 for the organic buffer (B) at the same flow rate for the analytical gradient. The gradient was as follows: 0 to 5% B over 10 minutes, 5% to 60% B over 100 minutes, then 60% to 100% B over 20 minutes. Over the course of this gradient fourteen 0.3 mL and twelve 0.5 mL fractions were collected, for a total of twenty-six fractions. Smaller fractions were collected for the first half of the gradient, where a more complex elution profile was observed. These fractions were stored at −20 °C prior to analysis.

**Figure 1:**
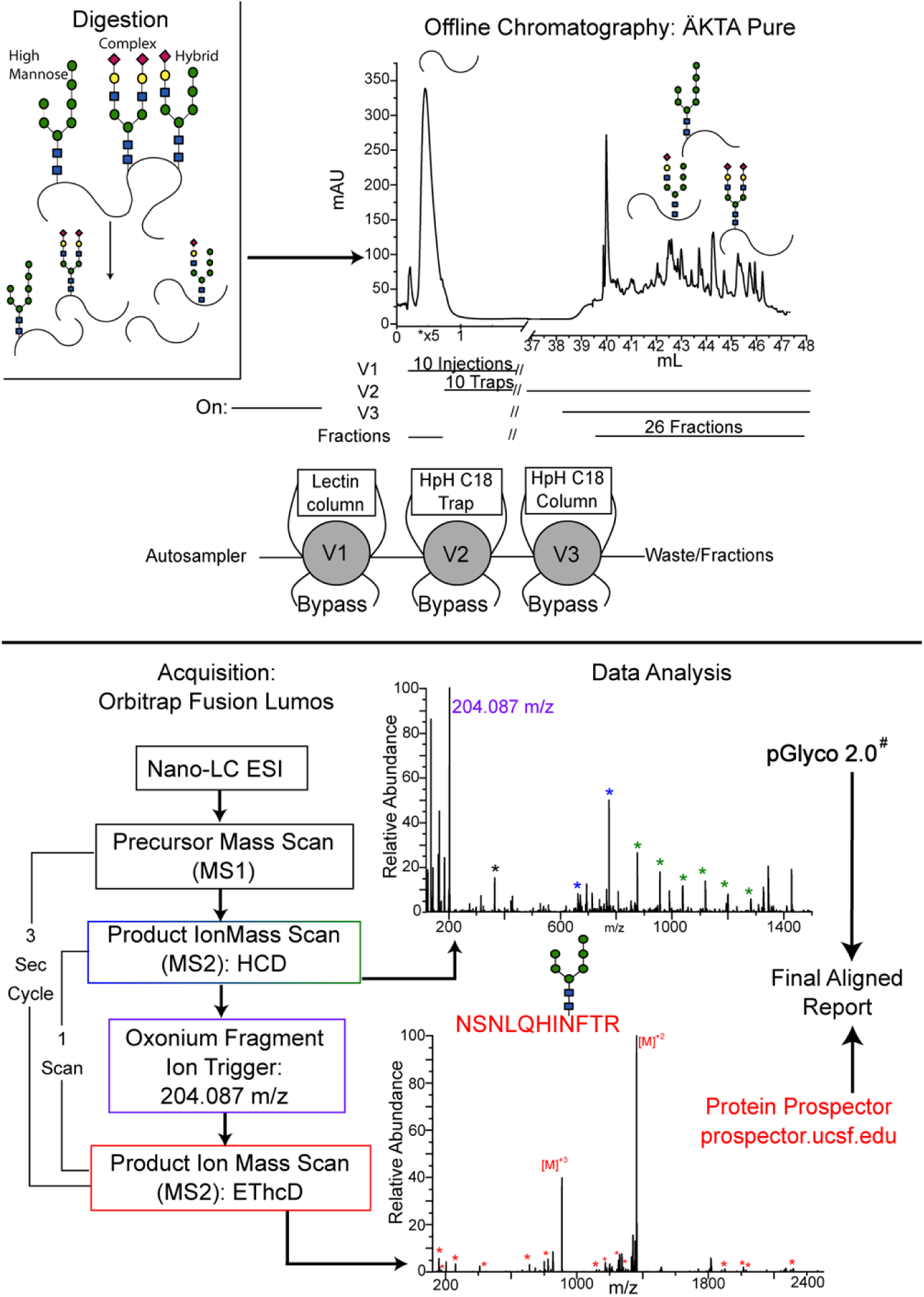
Experimental process diagram. Peptide digest is injected onto the three column ÄKTA Pure setup. An example final trap and fraction UV trace is shown, denoting where non-glycopeptides and glycopeptides are observed. The V# notations on the cartoon indicate the valves each of the three columns were placed on, while the lines below the UV trace indicate when each column was online. The LC-MS/MS acquisition parameters, along with an HCD spectra displaying diagnostic glycan fragments and the triggering oxonium ion are shown with an EThcD spectra displaying c and z ions used for fragmentation. The software used for data analysis with each method is noted. (Full width full page)

### LC-MS

Enriched peptides from the high *p*H reverse phase fractions were resolubilized in high purity water (ThermoFisher, Waltham, MA) with 0.1% formic acid (98.9% purity, ThermoFisher, Waltham, MA). Peptides were desalted (Acclaim PepMap 100, 75µm × 2 cm, nano viper, C18, 3 µm, 100 Å, ThermoFisher, Waltham, MA) and separated (Acclaim, PepMap RSLC, 75 µm × 25 cm, nanoviper, C18, 2 µm, 100 Å, ThermoFisher, Waltham, MA) using a Thermo Scientific Easy-nLC 1200 (ThermoFisher, Waltham, MA) over a 120 minute gradient at a flow rate of 300 nL/min. Buffer A was high purity water with 0.1% formic acid and buffer B was 80% acetonitrile in 19.9% high purity water with 0.1% formic acid. The gradient was as follows: 2 to 7% B over 30 seconds, 7 to 38% B over 100 minutes, 38% to 100% B over 10 minutes, then held for 9.5 minutes. Eluting glycopeptides were electrosprayed directly onto an ETD-enabled Thermo Fusion Lumos (ThermoFisher, Waltham, MA). Survey scans were performed at 60,000 resolving power in the Orbitrap mass analyzer with EasyIC internal recalibration. In addition to their use for EThcD-triggering, the HCD spectra were also used for glycopeptide identification using the software pGlyco (44). To promote the generation of y and b-type ions (in addition to Y and B-type glycosidic fragments), we conducted HCD at a relatively high, ramped collision energy (35% ± 5%) (44). This resulted in an overall higher number of HCD-identified glycopeptides, while still maintaining generation of the oxonimum ions used to trigger EThcD. Precursor selection occurred in the quadrupole with a 3 m/z isolation window, 0.5 m/z off set, and an AGC setting of 2.0 × 10^5^ or 200 ms before HCD fragment ion spectra were acquired. Ions with a charge state of 2 were selected if their mass range was between 750-2000, while ions with a charge state of 3-6 were selected if their mass was between 500-2000. Additional MS/MS spectra were acquired if HCD fragmentation generated an N-acetylglucosamine (GlcNAc, 204.0867, 15 ppm tolerance) diagnostic ion within the top 20 most abundant ions in the scan. The additional MS/MS scan involves selecting the same precursor ion and then subjecting it to EThcD fragmentation with a fluoranthene ETD reagent using the calibrated ETD reaction parameters, with supplemental HCD collision energy of 15, and an AGC setting of 1.5 × 10^5^ or 150 ms. All product ion scans were acquired in the Orbitrap mass analyzer at 30,000 resolving power and a mass range between 120-2500 m/z.

### Peptide Database Searching

Database searching was done using several software packages and online tools. Peaklists generated from HCD fragmentation were searched using pGlyco 2.1.2 (44, 45). Peaklists for EThcD data were generated by Proteome Discoverer 2.1.1.21 (ThermoFisher, Waltham, MA) and resulting spectra were submitted to Protein Prospector (46). Glycosylation lists used as variable modifications are denoted in Supplemental Table 2. Carbamidomethylation of cysteine residues was set as a fixed modification. A maximum of two missed tryptic cleavages were allowed. In both searches oxidation of methionine, acetylation of the N-terminus, pyroglutamate conversion of glutamate, and loss of protein N-terminal methionine were allowed as variable modifications with up to three total modifications per peptide. Precursor and product ion mass accuracy tolerance were set to 10 ppm. For both software packages glycosylation was only allowed to occur at asparagine with the N-X-S/T motif. Both datasets were searched against the Swiss-Prot human proteome, which contains 20,240 entries (downloaded November 2017). Randomized versions of these databases were generated by the respective software.

### Data Thresholding and Filtering

Peptide identifications from both searches were filtered to remove non-glycopeptide and probable false identifications. For Protein Prospector, glycopeptides were only accepted if the best scoring glycopeptide for that protein had an expectation value of 1×10^−6^ or less. For individual glycopeptides meeting these criteria, they also needed a peptide-level expectation value of 0.05 or less and a peptide score of 15 or greater. As an additional control, the data were also searched allowing for one non-tryptic cleavage (data not reported). Fully-tryptic results were then compared against this list to see if any semi-tryptic peptides produced better scores for a given spectra, and if so the data was manually inspected and the fully-tryptic entry removed if necessary. For the entire dataset, 5,445 unique forward database glycopeptides and 26 decoy glycopeptides were identified for a final EThcD FDR of 0.5%. For pGlyco, peptide and glycan scores are calculated independently. We initially required both peptide and glycan scores to be 10 or above. Duplicate identifications were then removed, with the highest overall scoring identification being kept. This resulted in 5 decoy identifications and 4,986 unique forward identifications for the entire dataset, for an HCD FDR of 0.1%. The appropriateness of individual score thresholds was confirmed by manually examining a set of low scoring glycopeptides.

In the cases of high mass, low intensity glycopeptides, we observed many instances where the monoisotopic peaks are either improperly assigned or entirely absent for the raw precursor scan. The resulting peaklists misidentified the precursor as approximately 1 Da heavier. This caused artifactual misidentifications in instances where two glycans in the database differed by 1 Da (47). This was often observed for two fucose subunits (292 Da) in place of a single sialic acid (291 Da). For instances where the same peptide was identified twice with glycans differing by 1 Da, the heavier identification was removed from the data if the observed retention time was within three minutes of the lighter glycan and the species occurred within three high *p*H RP fractions. This analysis was performed as the last step prior to calculating FDR on only the forward database identifications. The .Raw files, centroided peaklists, pGlyco annotated peaklists and Protein Prospector MS Viewer results have been uploaded to massive.ucsd.edu as part of the Proteome Xchange consortium under the dataset ID PXD013715 and can be accessed via ftp at ftp://MSV000083745@massive.ucsd.edu using the password “test_access”.

### Data analysis and Cytoscape

Identifications from Protein Prospector and pGlyco were aligned using Microsoft Access if the peptide, glycan, and other variable modification(s) matched. If identifications from Protein Prospector with an ambiguous glycan or modification position aligned with an unambiguous identification from pGlyco, the pGlyco identification was used for alignment and the Protein Prospector identification was no longer considered ambiguous. The identifications, both aligned and nonaligned, for each tissue type are reported in Supplemental Tables 3 and 4. Data analysis was preformed using Microsoft Excel and Origin Pro 2018. The most recent builds of Peptide Atlas (48) were downloaded for brain and serum and used without modification. Amino acid frequency data were acquired for all proteins in the Swiss-Prot human proteome (downloaded January 2019). R Studio and the R environment was used for Pearson pairwise correlations of glycosylation patterns. Cytoscape (49) was used for the network analysis and data visualization. AllegroLayout was used to generate the final layout of the glycan correlation network using the following parameters: 2,000 interactions, no overlapping interactions, independent component processing, component sorting, the Allegro Spring-Electric layout algorithm, 36% scale tuning, 100% rectangular gravity, with normalized edge weighting based off the R calculated and filtered correlation coefficients.

### Experimental Design and Statistical Rationale

One five mg sample of brain and one 10 mg sample of sera was analyzed. Each sample was glycopeptide enriched and fractionated into 26 high pH reverse phase fractions that were each in turn analyzed with two hour LC-MS/MS runs. Prior to the final analysis, the lectin enrichment and high pH reverse phase fraction steps were extensively tested to confirm reproducibility (data not shown). These experiments were aimed at providing an initial global perspective on human glycosylation and while technical replicates would provide additional glycopeptides due to the stochastic nature of data-dependent acquisition, acquiring an additional 104 hours of LC analysis was not deemed critical for an approximately 25 percent increase in total identifications. The robustness of the workflow can be observed from the successful completion of two distinct samples.

## Results

### Multi-lectin weak affinity chromatography development

Similar to other PTMs, N-linked glycopeptides are of low abundance in complex mixtures. As a consequence, many groups have developed methods which enrich glycopeptides from the large excess of non-modified peptides present in digests of complex samples (50). Adding to previous work, we sought to extend our ability to profile glycopeptides across a wide dynamic range through the optimization of multiple factors during development of the M-LWAC protocol presented here. These included efforts to minimize carryover of glycopeptides, maximize enrichment, and minimize sample loss. As our goal was to identify a maximum number of glycopeptides, we decreased individual sample complexity through the incorporation of high *p*H RP fractionation prior to LC/MS analysis. We designed an integrated setup that allowed us to reduce sample handling and processing steps between enrichment and fractionation, decreasing sample loss and increasing our overall yield (Figure 1). This consisted of a three-column system, which included: the M-LWAC column, a high-*p*H reverse phase (RP) trap column, and a high-*p*H RP analytical column. During the lectin chromatography phase, the trap column is switched inline to capture the glycopeptide eluent. This trapping step allowed use of optimized M-LWAC column flowrates in the absence of the analytical column (which could not be run at these higher flow rates). Our current approach utilizes a column that is eight percent the length our previous study, allowing for higher flowrates and minimizing carryover (42). Further details on the method development are provided in the Supplemental Materials and Methods.

### Glycopeptide mass spectrometry based acquisition

Recent advances in multi-mode fragmentation allow the acquisition of complementary fragmentation spectra on the time-scale of eluting peptide species.(51) The utility of this has been the focus of several recent studies (50–67). We acquired the data using an HCD-triggered EThcD approach relying on detection of an N-acetylglucosamine oxonium ion (4, 57, 68–74). To increase our specificity for selecting glycosylated ions for MS/MS, we compared the mass distribution of glycosylated and non-glycosylated peptides (SI Figure 1) and set our acquisition method to only select those species with precursor masses greater than 1,500 Da.

### HCD and EThcD provide complementary information

The HCD spectra were analyzed using pGlyco, which has the ability to search for glycosidic fragments as well as fragments occurring along the peptide backbone (44). EThcD spectra were searched using Protein Prospector (46), which has the ability to search these spectra for c, z, b, and y ions which retain the intact glycan. A given glycopeptide was positively identified using both types of fragmentation 39% of the time (2,928 out of the 7,503 instances). In some instances, this dual fragmentation approach allowed us to use HCD information to clarify partially ambiguous EThCD peptide spectral matches. One type of scenario we encountered dealt with a peptide sequence either modified with HexNAc2Hex9 or HexNAc2Hex8Fuc plus an oxidized methionine residue (SI Figure 2). In these cases, HCD could be used to determine the likely glycan composition.

**Figure 2:**
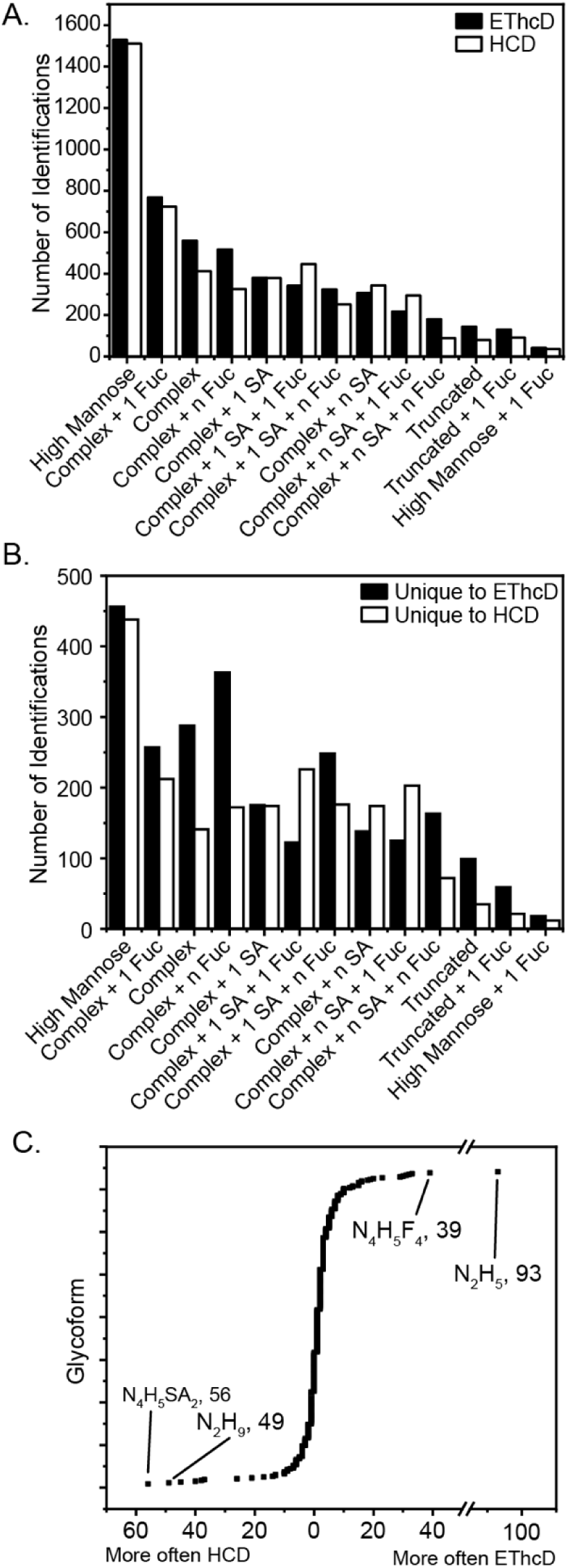
A) Glycoform family identifications from EThcD (black) or HCD (white) spectra observed in both human brain and serum glycoproteome. B) Glycoform family identifications for those glycopeptides identified by a single fragmentation mode, either EThcD (black) or HCD (white) C) The difference in the number of occurrences for individual glycoforms found by HCD versus EThcD, respectively from left to right. (two column, quarter page)

Because every glycan identified had a paired HCD and EThcD spectra, we were able to determine the relative success rate for each fragmentation mode (Figure 2). Overall, EThcD spectra (n = 5,445 glycopeptides) identified more glycopeptides than HCD spectra (n = 4,986 glycopeptides). To examine if these modes showed relative bias towards certain glycans, glycopeptides were grouped into truncated, high-mannose and complex categories which were further divided based on the number of fucose and sialic acid units. We find only subtle differences between fragmentation modes in the overall number of identifications for each glycan family, supporting the conclusion that both fragmentation modes are adequate at identifying glycopeptides with an array of glycoforms attached (Figure 2A).

Nevertheless, 4,575 glycopeptides were identified using only a single mode, and of these 55% (2,517) were identified from EThcD spectra while 45% (2,058) were from HCD spectra. When analyzing this subset of glycopeptides, we observed greater fragmentation specific differences (Figure 2B). Glycans belonging to the families including complex, complex with multiple fucosylation and/or multiple sialylation showed a greater number of identifications when using EThcD spectra. In contrast, singly fucosylated complex or multiply sialylated complex glycoforms were more readily sequenced when using HCD spectra.

Sialic acid will typically reduce the charge on multiply sialylated glycopeptides. Since EThcD fragmentation efficiency increases with increasing precursor charge state, multiple sialic acids would be expected to lower the relative success rate of EThcD relative to HCD. For sialylated complex glycopeptides, those with charge states less than 4 were much more likely to be identified by HCD. Furthermore, ∼80% of the EThcD spectra but only ∼40% of the HCD spectra identified this glycoform family with charge states 4 or higher. We the plotted the distribution of those glycoforms that are more frequently identified using a single fragmentation mode (Figure 2C). This data reinforces the observations mentioned above, with the multiply sialylated complex glycoform (HexNAc_4_Hex_5_NeuAc_2_) being identified 56 more times from the HCD spectra compared to EThcD. In contrast, the multiply fucosylated complex glycoform HexNAc_4_Hex_5_Fuc_4_ was identified 39 more times in EThcD spectra compared to HCD spectra.

### Tissue-specific differences in the N-linked glycoproteome

We identified 666 glycoproteins in the entire dataset, with 192 proteins overlapping between both tissue types, 48 proteins unique to sera, and 426 proteins unique to brain (Figure 3A). It is important to note that the isolated brain tissue contained vasculature, and therefore proteins identified in both sera and brain tissue could nevertheless be blood-specific and not necessarily expressed by neurons or their supporting cells.

**Figure 3.**
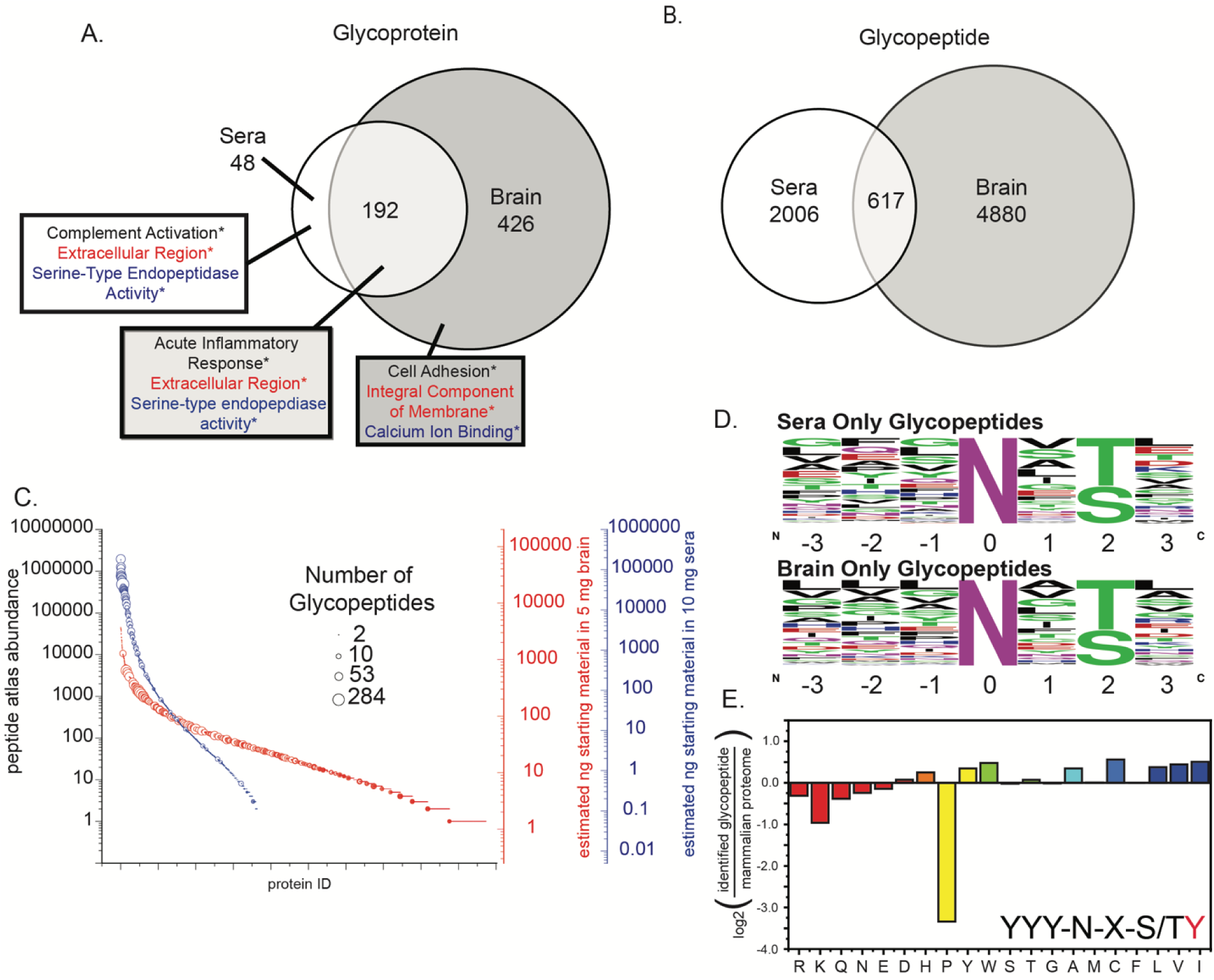
A) Number of unique glycoproteins observed in each tissue type. Gene ontology analysis for each of these groups of identified proteins showing the highest enriched biological process, cellular component and molecular function, respectively as black, red, and blue. Enrichment level determined using a background proteome of the top 250 most abundant proteins observed in Peptide Atlas for both brain and sera. Enrichment level p-value < 0.005 for each ID. B) Number of unique glycopeptides observed in each tissue type. C. Number of unique glycopeptides mapped (point size) onto proteins observed in Peptide Atlas analysis ranked by abundance for both sera (blue) and brain (red) tissue. Estimation of abundance (ng) found in starting material before enrichment using multi-LWAC. Proteins not found in glycoproteome are plotted as point with zero internal area. D. Amino acid frequency in sequence surrounding the site of glycosylation for unique peptides in each tissue type. E. Amino acid fold change at site +3 from the site of glycosylation relative to the natural abundance in the mammalian proteome. Hydrophobicity score is mapped to the fill color from hydrophilic (red) to hydrophobic (blue). (two column, half page)

Next, we examined the enrichment of gene ontology terms for proteins modified by each glycan family (75, 76). For this analysis, we used a protein background consisting of the top 250 proteins listed in Peptide Atlas both in brain or serum (48). Of our glycoproteins, 96% (632) were previously annotated as glycosylated (p < 4.3 × 10^−282^). Proteins found in serum showed an enrichment for the term extracellular region. We found that 33 of the 48 proteins unique to serum (p < 2.7 × 10 ^−17^) and 102 of the 188 proteins found in both serum and brain (p < 1.1 × 10 ^−37^) mapped to this gene ontology term. Proteins found only glycosylated in brain tissue were enriched for being integral components of the membrane (i.e., transmembrane domain, 310/425, p < 1.8 × 10 ^−64^). For the case of glycoproteins identified in both tissues, the enrichment for the term “extracellular region” is most likely driven by the serum component present in the brain vascular system.

We plotted the abundance of identified glycoproteins using the data in the Peptide Atlas Project (48). Figure 3C shows the dynamic range of the proteins in each tissue type, with the size of the point representing the number of glycopeptides identified for that protein. Furthermore, we have approximated the total mass of each protein in the starting material, reported on the right axis. This approximation is based off of the spectral counts acquired in the Peptide Atlas project and how they scale to the total protein starting material utilized in this analysis. We estimate that our enrichment allows us to measure glycoproteins over at least 4-orders of magnitude. For proteins annotated as glycosylated, we can gain further information on the effect that the dynamic range of a tissue’s protein abundance has on the glycopeptide identification success rate.

Figure 4 shows the identification rate of glycoproteins at different binned abundance windows, derived from Peptide Atlas. It is immediately obvious that in serum we have the majority of our identifications coming from the more abundant protein bins, which is followed by a rapid identification rate decay with lower abundance bins. In brain, however, the glycoprotein identification percentage decays more gradually as a function of decreasing protein abundance. These observations, when taken together, illustrate the challenges of working with serum derived protein mixtures. The top twenty proteins make up approximately 99% of the mass in sera (77), leading to lower abundance proteins being in the ng/mL to sub-ng/mL regime. This data demonstrates that the distinct abundance distribution between serum and brain has a significant impact on identifications, as a much larger number of brain glycoproteins were identified despite loading identical total protein amounts in both analyses.

**Figure 4.**
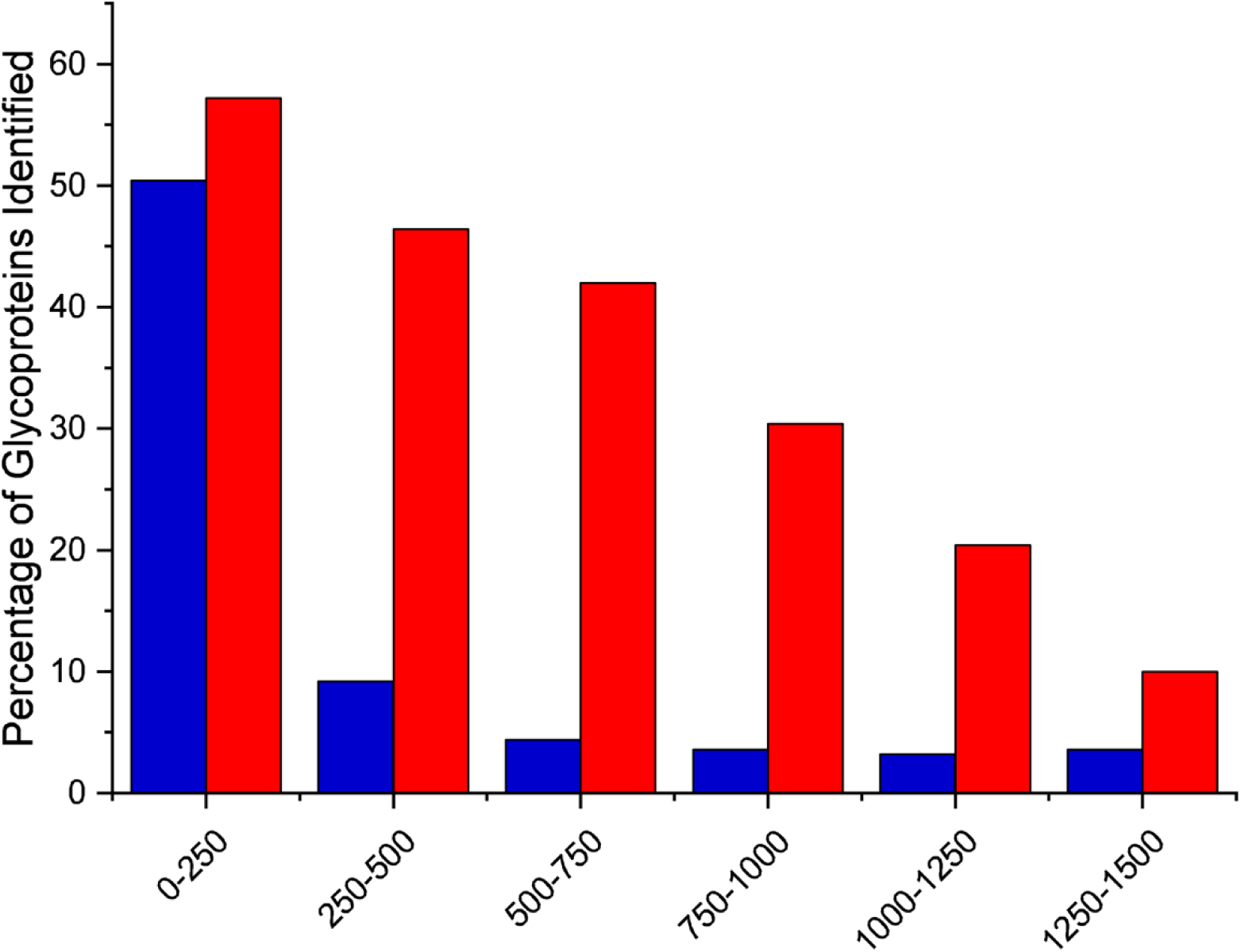
Success rate of glycoprotein identification as a function of Peptide Alas derived binned abundance for sera and brain proteome, blue and red respectively. Those proteins included were annotated in UniProt as being glycosylated. (one column, quarter page)

Our analysis identified 7,503 glycopeptides for the two tissue types, with 617 occurring in both datasets (Figure 3B). There were 2,006 glycopeptides identified in serum alone, and 4,880 glycopeptides identified in the brain tissue alone. We first examined the frequency of each amino acid residue adjacent to the N-X-S/T modification site (Figure 3D). We note, similar to Mann et al. (78), a high frequency of non-polar amino acid residues found to the N-terminal side of the PTM site in both tissues. For the two tissue types we noted no significant difference between the percentage of threonine (∼60%) and serine (∼40%) at position +2. We next calculated the enrichment of each amino acid residue surrounding the N-X-S/T compared to that amino acids abundance in the human proteome. When analyzing the enrichment of amino acids in more detail (SI Figure 3) we found, consistent with Mann et al. (78), that the +3 position had a much lower probability of proline (2,177 fold lower) (Figure 3E). We also observed that hydrophilic amino acid residues were less common at the –2, −1, +1 and +3 positions, while hydrophobic residues were more common at these same positions. This effect was most prominent at the +1 site, followed closely by the +3 site.

We next chose to take a glycan-centric view of the data, highlighting the differences between the tissue types. To analyze these differences, we calculated the percentage of uniquely identified glycopeptides with a glycoform belonging to one of the previously mentioned thirteen glycan families, defined above, for both serum and brain (Figure 5A, blue and red, respectively). We first noticed a discrepancy in the abundance of high mannose between the two tissue types. High mannose represents the largest glycan family in brain, comprising ∼35% (2,060 glycopeptides) of brain glycopeptide identifications, while it is only the third most identified in serum at ∼15% (407 glycopeptides) of identifications. Since the brain tissue is a whole cell lysate compared to the “secreted” serum proteome, it seemed likely that this difference is due to the higher amount of membrane content in brain compared to sera. Gene Ontology analysis confirmed that the major annotation for high mannose expressing proteins was “intrinsic to membrane” (73.4%, p < 7.9 × 10^−16^). When investigating high mannose modified proteins present only in sera, “intrinsic to membrane” was annotated for 68.6% of them (p < 8.9 × 10^−2^) and this was that highest percentage term for this set of proteins.

**Figure 5:**
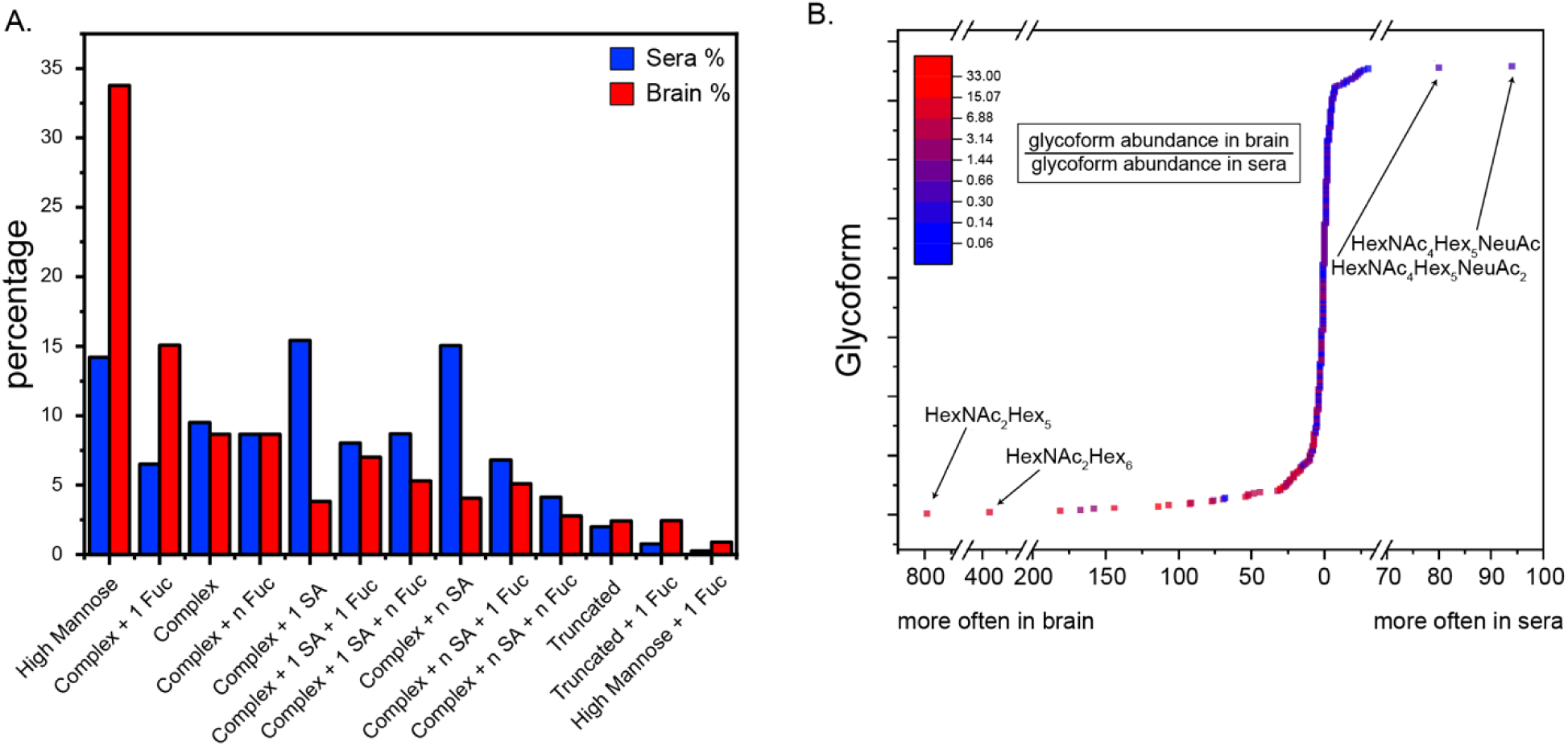
A) The glycoform family abundance is provided as an inner-tissue normalized percentage for sera (blue) or brain (red). B) The difference in individual glycoform identifications is given showing glycoforms more abundant in brain or sera, from left to right. The corresponding fold increase in brain relative to sera is shown color mapped. For example, HexNAc2Hex5 is 13.4-fold more abundant in brain than in sera. (1.5 column, quarter page)

Glycans were found to be sialylated at a higher frequency in sera (>57%; 1,614/2,876), compared to the brain proteome (<27%; 1,596/5836). There was no apparent enrichment for specific Gene Ontology terms when analyzing the sialylated glycoproteins, either overall or for the tissue-specific subsets. The set of proteins modified by complex glycans lacking sialyation also showed no significant enrichment of Gene Ontology terms. When the entire set of complex glycans (with or without sialyation) was examined together, the following terms were identified as enriched: “extracellular region” (9.4 fold enrichment, p< 6.5 × 10^−12^), “complement and coagulation cascade” (2.2 fold enrichment, p < 8.3 × 10^−6^) and “endopeptidase activity” (2.1 fold enrichment, p < 1.2 × 10^−4^).

The abundance of individual glycans within each tissue type was analyzed to examine differences based on exact glycan composition. In serum, the five most common glycans are all sialylated, with three being doubly sialylated. While prevalent, these three multiply sialylated glycoforms only account approximately 10% of the total sera identifications. By contrast four of the five most common glycans in brain were of the high mannose type, and these species account for almost 30% of all identifications in brain. Of those four, 20% of the total identifications in brain come solely from HexNAc_2_Hex_5_ and HexNAc_2_Hex_6_. These two correspond to the smallest mature high mannose glycans. The truncated glycans, HexNAc_2_Hex_4_ and smaller, constitute 6% of the identified glycopeptides in brain. The low overall abundance of truncated species suggests that post-mortem glycosidase activity is not a major factor influencing our observed glycopeptide distribution. We analyzed the relative fold-difference of individual glycans between the brain and serum (Figure 5B, color mapped). Although highly abundant glycans were found in some instances to display large changes in tissue abundance, those glycans with the greatest differential expression were not the most abundant.

### Regulation of glycan microheterogenity

We then investigated microheterogeneity at the glycopeptide level for the entire dataset. Figure 6 is a histogram plotting how many peptides bore a given number of different glycans. The majority peptides are identified bearing one to three glycans, with the average number of glycans per site equal to 4.07 and the median number of glycans per site equal to two. Approximately 10% of the identified peptides were modified by 10 or more glycans. These sites which displayed a large degree of microheterogeneity did not contain any obvious primary amino acid structural motif (data not shown), indicating that other protein structural elements likely drive the observed microheterogeneity differences.

**Figure 6:**
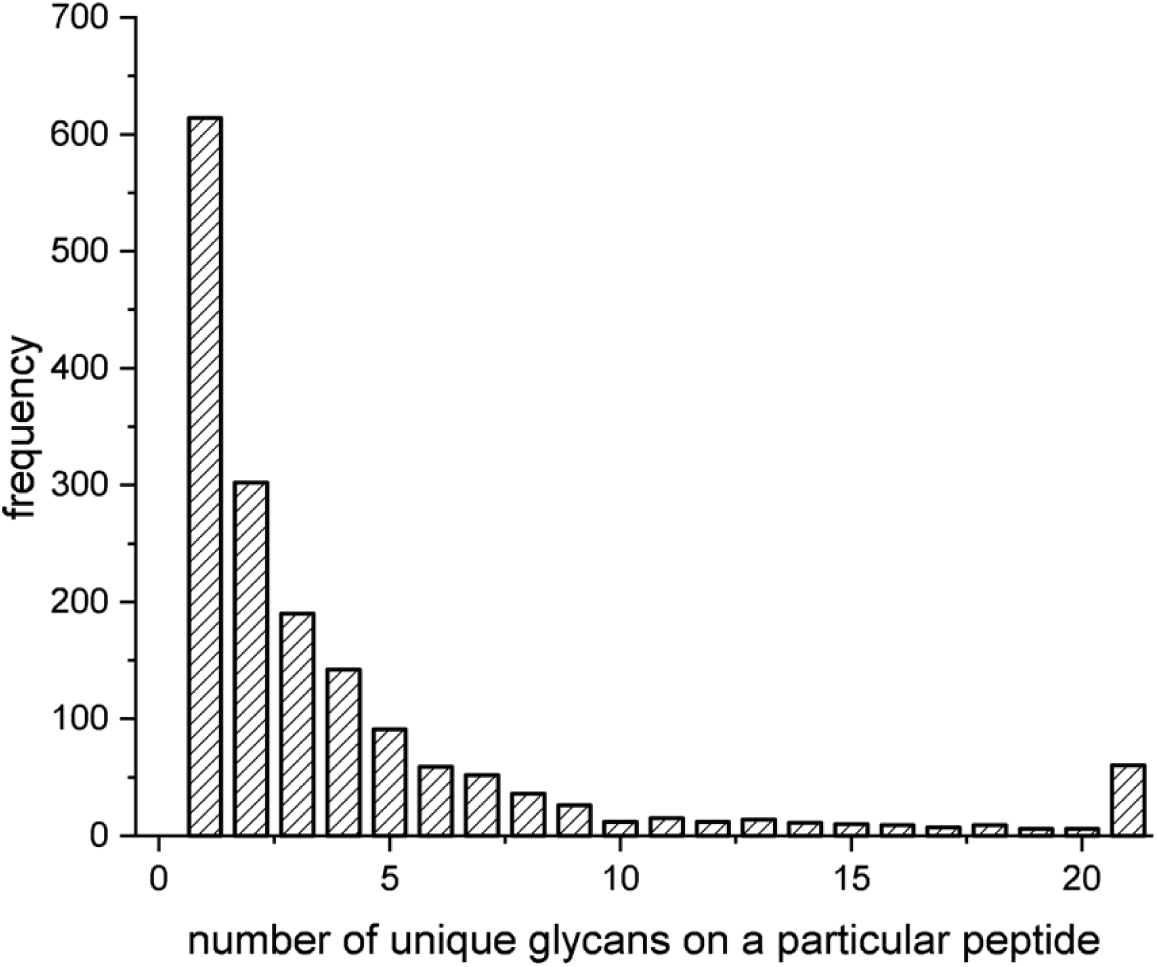
Glycopeptide microheterogenety is shown as binned groups of numbers of peptides with a particular number of observed glycoforms attached to the peptide. Peptides which are modified by more than 20 glycoforms are binned together. (one column, quarter page)

To understand how site microheterogenity may be regulated at the protein level, we investigated proteins for which at least two unique glycosylation sites were identified. A high number of glycans identified at one site correlated with a higher average number of glycans at the remaining sites (Figure 7). To control for bias based on abundance effects of glycopeptide detection, we also examined the subset of high-abundance proteins and the same trend was found within this group (data not shown). This provides evidence regarding the degree to which protein structure drives microheterogenity levels.

**Figure 7:**
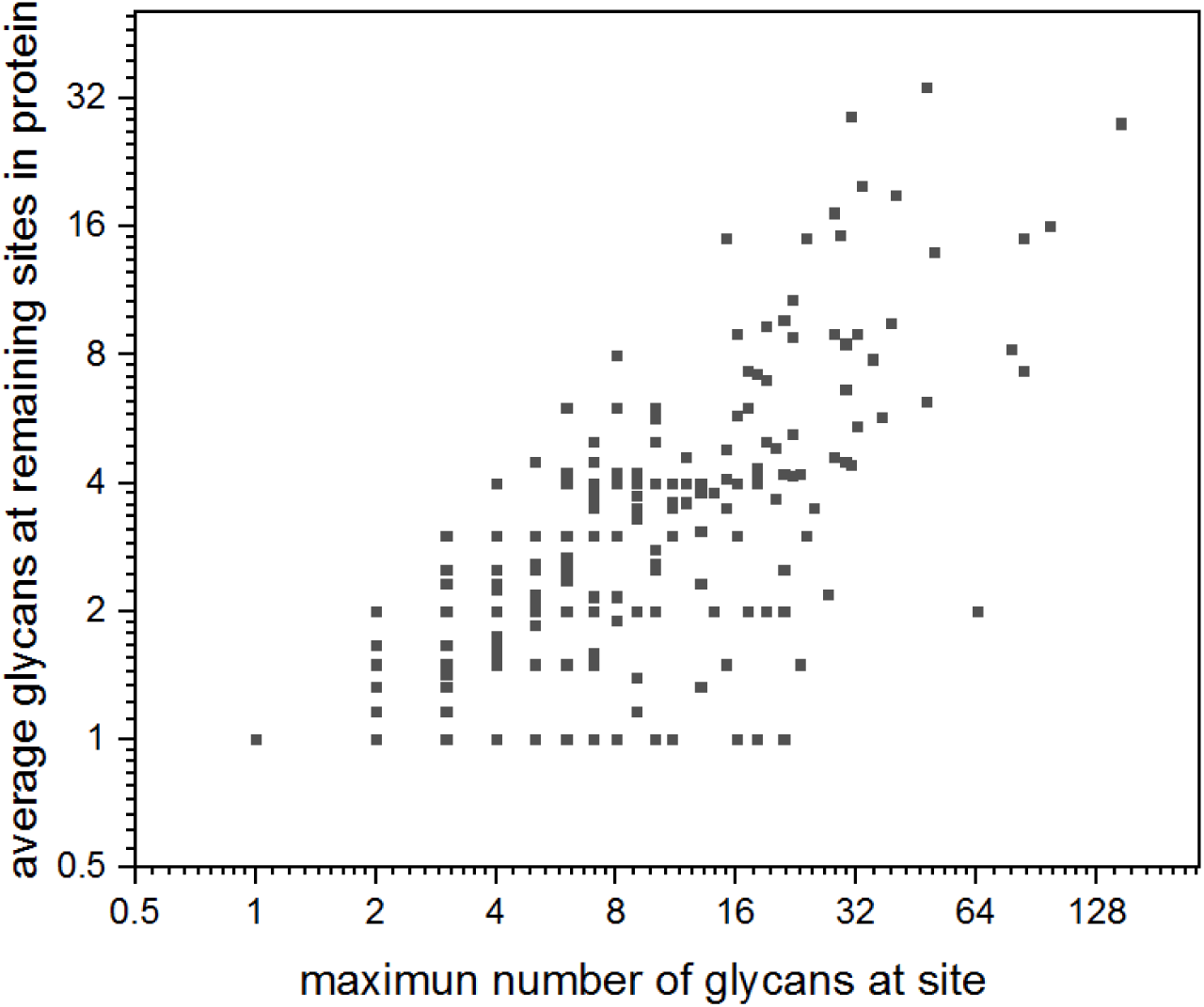
The maximum number of glycoforms found at any site on a protein is shown versus the average number of glycoforms per site for the remaining sites on the same protein. Proteins with greater than one identified glycosite are included in the plot below. (one column, quarter page)

We monitored the number of unique glycoforms originating from a specific protein versus the number of identified sites of glycosylation, as a metric to determine particular proteins which are more diversely glycosylated (Figure 8). The average number of glycans per site was roughly 2.5 times greater in serum (6.94 glycans/sites, Figure 8, blue line) compared to that within brain (2.70 glycans/site, Figure 8, red line). This effect may be due to the overall accessibility of transmembrane versus secreted proteins.

**Figure 8:**
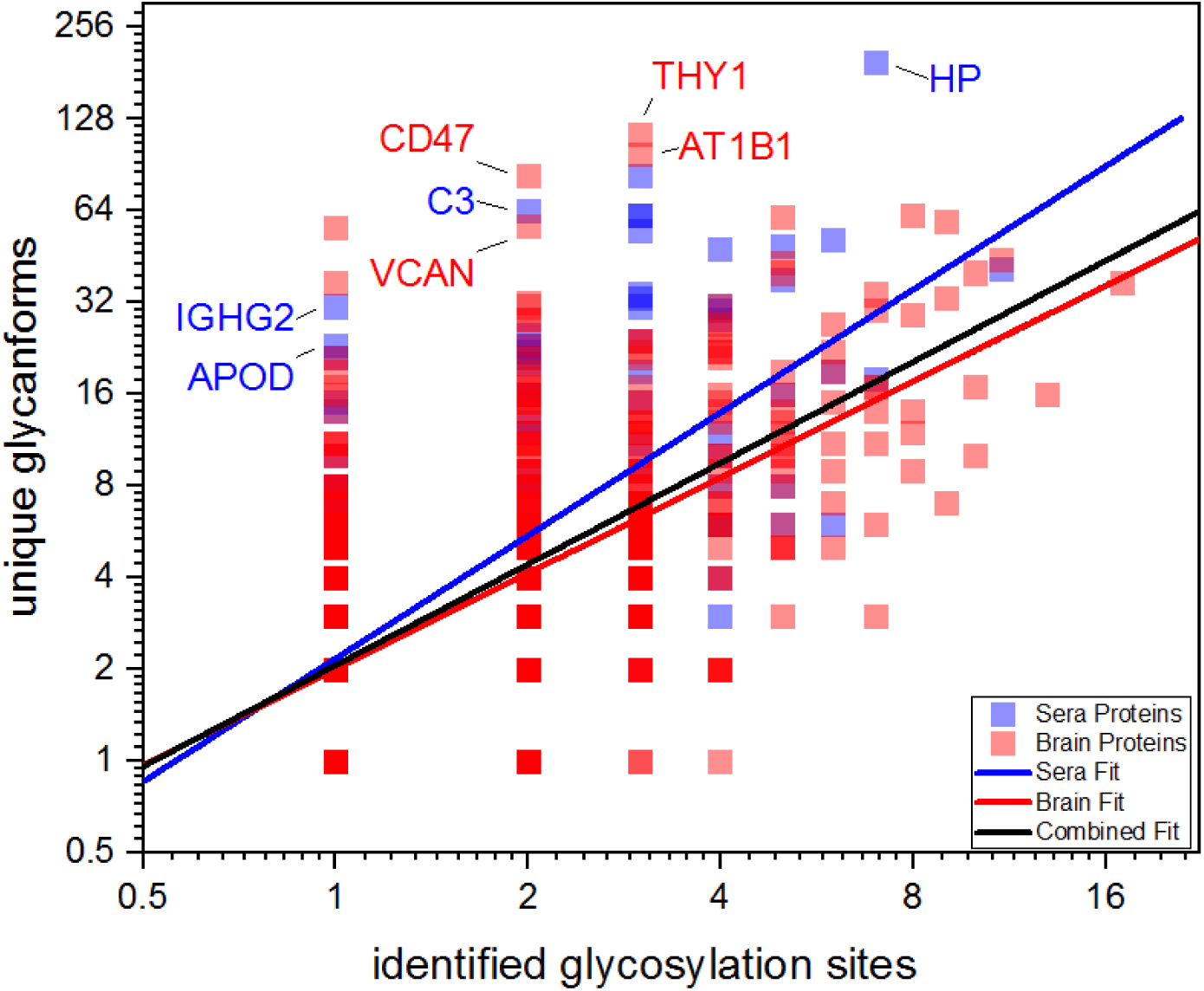
The number of identified glycosylation sites is plotted against the number of unique glycans found attached to each protein found in brain sera (blue) or brain (red). The linear fits represent the fitted number of glycans per sites of glycosylation are given for sera (slope = 6.94, blue), brain (slope = 2.70, red) and the combined datasets (slope = 3.39, black). Individual gene names are provided for the genes with the largest ratios of unique glycans per sites of glycosylation. (one column, quarter page)

### Glycosylation processing networks

To gain insight into potential differences in glycan biosynthesis, we examined the canonical glycan processing pathway involving Man_9_GlcNAc_2_ down to Man_5_GlcNAc_2_ and the subsequent initial steps in the synthesis of complex glycans (Figure 9A). Each glycan is represented by a short name, a drawing of its possible branching pattern, and a corresponding node in both the serum and brain datasets. Each node size is proportional to the number glycopeptides identified bearing that glycan, while the thickness of each connecting edge is proportional to the frequency of modification co-occurrence. Co-occurrence percentage was calculated by determining the total number of unique peptides found modified with both glycans divided by the number of peptides found modified with either glycan. Numerous differences can be observed between tissue types. For example, there is an inverted order of glycan abundances for the high mannose glycoforms within each tissue, with the initial steps most abundant in sera and the Man_5_GlcNAc_2_ end most abundant in brain. In both tissues, there is a marked decline in abundance between Man_5_GlcNAc_2_ and Man_5_GlcNAc_3_, indicating that MGAT1 activity (which transfers GlcNAc to one of the arms) is one of the rate limiting steps in complex N-glycan synthesis in these tissues. The overall co-occurrence of high mannose species within the brain is higher than in serum, as is indicated by thicker lines connecting individual high mannose nodes. For the addition of core fucosylation, there are notable differences both between and within tissues. In sera, Man3GlcNAc4 is well correlated with Man_3_GlcNAc_4_Fuc, while in brain, the strongest fucosylation correlation is between Man_3_GlcNAc_5_ and Man_3_GlcNAc_5_Fuc. In contrast, we detected no peptides in sera that had versions modified with Man_3_GlcNAc_3_ and Man_3_GlcNAc_3_Fuc.

**Figure 9:**
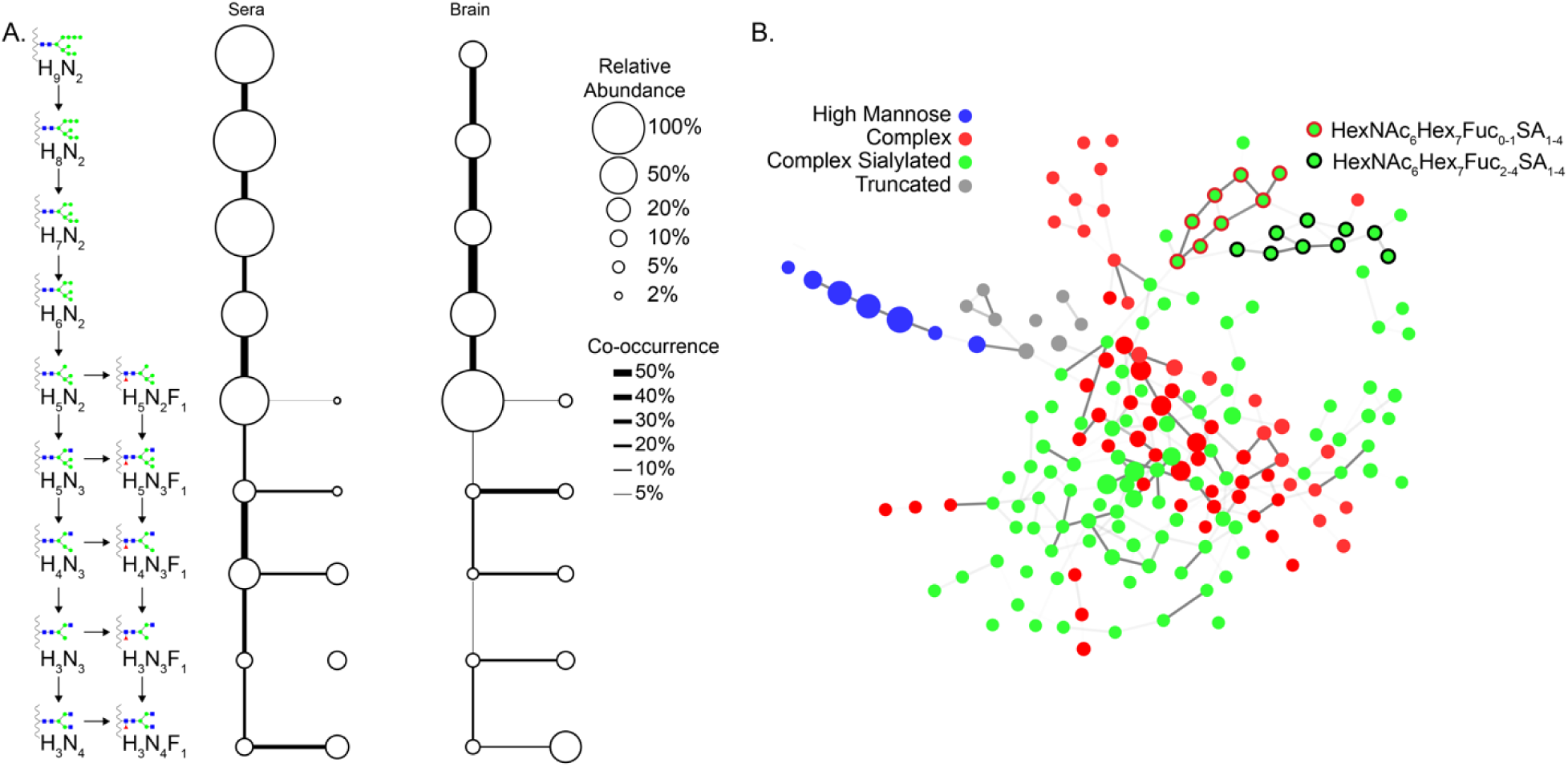
A. The co-occurrence rate for the specific glycans (left) found along the common pathway of N-linked glycan biogenesis. Size of circles represent the number of a particular glycoform identified (normalized within tissue type). Line thickness represents the level of co-occurrence between adjacent glycoforms (thicker lines represent greater factor of co-occurrences). Common fucosylated subgroups are provided as horizontal. B. Glycan co-occurrence networks were generated for all glycans, where the number of identified glycopeptides with a particular glycan scaled the size of each node. Different families of glycans were individually color coded. Edges connecting nodes were scaled to Pearson correlation coefficients of glycoforms on peptide identifications. Individual glycan groups were identified as forming subnetworks, including HexNAc_6_Hex_7_Fuc_0-1_ and HexNAc_6_Hex_7_Fuc_2-4_SA_1-4_. Cytoscape application AllegroLayout was used to gravity orient the network. (two column, half page)

To create a network view of interconnection between glycans, we plotted each glycan as a node proportional to the number of observations of that glycoform in the entire tissue dataset (Figure 9B). Co-occurrence between glycans was calculated using Pearson pairwise correlation coefficients. We only considered edges between glycans that differed by one monosaccharide unit, which theoretically can be directly converted via enzymatic activity. Only edges with correlation coefficients equal or greater than 0.22 are shown, for a total of 300 edges. These edges are shown with different degrees of transparency based off of increasing correlation coefficient. Following this, we used the Allegro Layout in Cytoscape to self-organize our network, effectively grouping nodes of higher correlating glycans closer together (49).

High mannose glycans formed a cluster which was well separated from the overall population (blue, Figure 9B). Complex non-sialylated glycoforms (red) were partially isolated from complex sialylated glycoforms (green). This unique patterning could be an outcome of a highly correlating core group of complex glycoforms which are not equally correlated with their surrounding sialylated members. Finally, we note two more tailored groupings, specifically the HexNAc_6_Hex_7_Fuc_0-1_SA_1-4_ (green nodes, red outline), and the HexNAc_6_Hex_7_Fuc_2-4_SA_1-4_ (green nodes, black outline). These nodes are strongly connected by edges within their subgroupings, indicating a high degree of co-occurrence within this region of the glycosylation processing pathway.

## Discussion

### Considerations for enrichment and acquisition

While our enrichment strategy was effective, the optimal approach for glycopeptide analysis will depend on sample specific factors such as: tissue, organism, initial total protein mass, sample complexity and glycoforms of interest. Our workflow involves a relatively complex HPLC setup which includes an auto sampler, three columns, and a fraction collector. As such, this approach may only be beneficial for highly complex samples. We find that the M-LWAC and the high *p*H reverse phase trap are effective for isolating glycopeptides in a single enrichment, and can be directly analyzed by LC/MS without additional multidimensional fractionation. In regards to the complexity of the sample, fractionation will increase the total number of identifications, but does not necessarily need to be coupled directly to the lectin enrichment step (31, 79). Our goal in coupling these steps was to minimize the overall sample handling, as it correlates with increased inter-sample variation and decreased yield. A comparison of M-LWAC with fractionation versus without fractionation, shows that identifications do not scale linearly with starting amounts or number of fractionations. We found that enrichment without fractionation of 100 µg of whole brain lysate produced approximately 400 glycopeptide identifications in a single LC/MS analysis. In contrast, with 50-fold more sample and high pH fractionation, we identified approximately 12-fold more glycopeptides.

Enrichment using lectins likely plays a large role in the exact species identified. One common alternative enrichment method is hydrophilic interaction liquid chromatography (HILIC), which utilizes the higher hydrophilic nature of glycopeptides compared to unmodified peptides (32, 64, 70, 80). While HILIC relies on the increased hydrophilicity of glycopeptides compared to other peptides, lectins have specificity for mono or disaccharide residues. Utilizing a single lectin can therefore be used for targeted glycoform enrichment, which would be desirable for profiling specific changes in disease states (10, 12, 13, 81, 82). HILIC may provide a broader enrichment than a single lectin approach. However, during HILIC, glycopeptides bearing polar sialic acid residues bind much more effectively than non-sialylated glycopeptides (83).

The choice of gas phase fragmentation mode is an important factor for glycopeptide analysis. The specific fragmentation mode controls the number of spectra acquired as well as the available search engines used for analysis of generated fragment ions. An EThcD scan can take 2 to 10 times as long as an HCD scan, significantly affecting the total number of species for which data is acquired. Our data demonstrates that EThcD outperforms HCD for a subset of glycoforms. This was especially apparent for complex and complex + n fucosylated glycoforms. We also found that higher energy HCD is required to obtain y and b ions from glycopeptides relative to non-glycosylated peptides. However, a balance needs to be struck between higher energies, which create optimal levels of y and b ions and lower energies which create optimal levels of glycosidic backbone Y and B ions (63) and the solution adopted by us and others is to employ a normalized HCD collision energy profile of 35 ± 5. Our overall dataset contained many examples of spectra which did not pass statistical threshold due in some instances to insufficient y and b ion levels, while other spectra demonstrated insufficient levels of Y and B ions. The latter case is likely due to “overfragmentation” of the spectra in an attempt to generate y and b ions. Future efforts will be aimed at developing HCD collision energies which account for peptide mass and charge.

For complex and the complex multiply fucosylated glycopeptides, we observed more unique identifications from EThcD generated spectra. We then examined the paired, low scoring HCD spectra from those glycopeptides to better understand why these spectra fell below the pGlyco score cutoffs. For confident identification of HCD spectra, the search engine pGlyco requires high confidence interpretation of both glycosidic fragments as well as fragments from the peptide backbone. For the majority of cases where the HCD spectra was of insufficient quality, the glycan score was below threshold, while the peptide score was sufficient. This same trend of low HCD glycan score was also identified for glycopeptides of the high mannose family that generated successful EThcD identifications. We hypothesize that this subset of HCD spectra that do not pass the glycan score threshold most likely fail due to too high of a collision energy. This high collision energy causes over-fragmentation of the glycopeptide, generating relatively insufficient amounts of glycosidic Y ion fragments. The low number of core ions identified by PGlyco, corresponding to the Y series ions, for non-passing HCD spectra identifications further supports that notion.

### Glycoproteome analysis: individual protein glyco-heterogeneity

Figure 8 provides evidence for the correlation of identified protein glycoforms with the number of identified protein glycosylation sites. In sera (Figure 8, blue symbols), we found apolipoprotein D (APOD) showed a very high level of macroheterogeneity, with 23 glycans at one site. N-linked glycosylation on this protein has been implicated as a biomarker in autisim spectrum disorder (84). The immune proteins complement C3 and immunoglobulin heavy constant gamma 2 (IGHG2) showed 64 glycans at two sites and 31 glycans at one site, respectively, both showing increased levels of macroheterogeneity. This level of diverse glycosylation may reflect the critical role glycosylation plays in the immune system (85). Finally, we found that the abundant blood protein, haptoglobin (HP) showed 196 glycans at seven sites. This protein is especially interesting as it has been implicated as a biomarker for many types of cancer, and represents the most diversely glycosylated protein we identified in this study (81, 86).

In brain tissue, the proteins hyraluronan and proteoglycan link protein 1 (HAPLN1, not shown) and versican core protein (VCAN) (87) were observed with 56 glycans at one site and 57 glycans at two sites, respectively (Figure 8, red symbols). These two proteins are implicated in a range of cancers, including the 29-fold upregulation in adenoid cystic carcinomas (88). Interestingly, these two proteins interact *in vivo*. Myelin-oligodendrocyte glycoprotein (not shown) was observed with 37 glycans at one sites. This protein is implicated in the pathogenesis of multiple sclerosis, specifically through the modulation of N-linked glycosylation and changes to the auto-immunogenicity of this protein (89). Additionally, the Na^+^/K^+^ transporting ATPase subunit beta-1 (ATP1B1) was identified with 98 glycans at three sites. While this protein functions as an ion transporting protein, it has also been observed to modify its N-glycan structure to control cell adhesion (90). Another cell-adhesion protein we identified is Thy-1 membrane glycoprotein with 114 glycans at three sites. This protein has been correlated with cell transformation in breast cancer tissue (91). The protein leukocyte surface antigen CD47 was observed with 83 glycans at two sites. N-linked glycosylation of CD47 regulates localization of this immune modulating protein and aberrant CD47 glycosylation on the surface of ovarian cancer cells allows enables immunological evasion (92).

As proteins are glycosylated through the endoplasmic reticulum and Golgi apparatus, a host of glycosyltransferases and glycosidases act to determine the final collection of glycan structures which ultimately modify a given site. As shown in Figure 9A, analyzing data at the intact glycopeptide level allows us to infer the efficiency between individual steps in the synthesis pathway. Furthermore, when this analysis is done without the canonical pathway restrictions, glycoform families are shown to cluster together (Figure 9B). This illustrates those subpopulations whose biosynthetic steps may be more tightly correlated. Functional characterization focusing on the site-specific co-occurrence (or relative levels) of related glycans can act as a novel metric to profile changes in physiological states and may act as a new avenue for biomarker discovery.

In summary, this dataset represents the largest human serum and the only brain intact glycoproteome to date. Careful optimization and implementation of a multi-lectin weak affinity chromatography platform along with high resolution mass spectrometry allowed the mapping of 379 glycoforms, to 7,503 unique glycopeptides, originating from 666 glycoproteins over a dynamic range of four orders of magnitude. Certain glycans displayed moderate preference for HCD versus EThcD fragmentation which likely stems from the effect of sialic acid on precursor charge state as well as how glycan length and branching affects search engine scoring. Glycopeptide microheterogeneity was widespread, with as many as 199 distinct glycans being identified at a specific site. Proteins with a high degree of microheterogenity at one site had an above-average degree of microheterogenity at other sites, indicating that this process is partially regulated at the protein level. Finally, we demonstrate that examining glycan co-occurrence can provide indirect information regarding the underlying biosynthetic pathways, with brain-derived glycopeptides bearing high-mannose glycans more likely to be identified with similar high-mannose glycans than corresponding sera-derived glycopeptides. This type of data analysis has potential application as a novel metric to measure glycan metabolic changes during disease.

## Acknowledgements

This work was supported by National Institutes of Health Grant number 5R01GM117207 (D.E.C), K01 MH107756 (M.L.M), and Indiana University Precision Health Grand Challenge Initiative (J.C.T).^1^

## Data Availability

The .Raw files, centroided peaklists, pGlyco annotated peaklists and Protein Prospector MS Viewer results have been uploaded to massive.ucsd.edu as part of the Proteome Xchange consortium under the dataset ID PXD013715 and can be accessed via ftp at ftp://MSV000083745@massive.ucsd.edu using the password “test_access”.

## Footnotes

1 The content is solely the responsibility of the authors and does not necessarily represent the official views of the National Institutes of Health.

